# Cytosolic fructose - an underestimated player in the regulation of the sucrose biosynthesis?

**DOI:** 10.1101/2024.12.11.628007

**Authors:** Oliver Giesbrecht, Christina Bonn, Lisa Fürtauer

## Abstract

Plants must continuously adapt to environmental fluctuations, which significantly influence their photosynthetic performance and overall metabolism. The sucrose cycling system within plant cells plays a critical regulatory role during stress conditions. This study employed a systems biology approach to analyze system stabilities mathematically under various regulatory conditions impacting sucrose cycling dynamics. We investigated the effects of mutations within this cycle, specifically HEXOKINASE1 (*Arabidopsis thaliana gin2-1*), alongside high-light exposure. Finally, we confirmed the modelling output in vitro by enzyme assays. The implementation of experimental subcellular metabolite data into a Structural Kinetic Model (SKM) enabled exploration of regulatory responses and system stabilities within a three-compartment model. Within system instabilities, *gin2-1* was more instable than its wild type. The *gin2-1* mutation particularly was destabilized when fructokinase function was impaired by phosphorylated sugars. Additionally, we confirmed that phosphorylated sugars serve as stronger activators of sucrose-phosphate synthase (SPS) than glucose does. Interestingly, models with fructose SPS activation exhibited a similar stability pattern. Consequently, we proposed and confirmed in silico a triple activation of SPS by highly activating phosphorylated sugars and lower activating nonphosphorylated hexoses. Additionally, we biochemically confirmed the previously unknown, but now predicted, activation of SPS by fructose in vitro. In summary, our study highlights the essential role of sucrose cycling in plant cells under stress conditions. The in silico findings reveal that phosphorylated sugars are stronger activators of SPS than glucose and introduce a previously unknown activation mechanism by fructose. These potential activation capacities were confirmed in vitro through SPS enzyme activity assays, underscoring the efficiency of our systems biology approach. Overall, this research provides valuable insights into carbohydrate metabolism regulation and paves the way for future investigations to deepen our understanding of the complexities involved in sucrose cycling and biosynthesis in plants.

## Introduction

Plants need to constantly adapt to their surrounding environmental fluctuations, which influence their photosynthetic performance, anabolism and catabolism as well as reproduction and survival. One significant aspect of these fluctuations is changes in light availability, as photosynthetic cells have to adapt to varied intensities across a wide range of scales for instance temporary shading (canopy or clouds), day-night cycles, and seasons. Changed light intensities have a direct impact within chloroplasts on photochemical reactions, the light harvesting, electron transport as well as on the formation of ATP and NADPH and their availability for the Calvin-Benson-Bassham (CBB) cycle for synthesis of sugars [1]. Consequently, CO_2_ assimilation rates are significantly influenced by different light intensities and fluctuations during a day or growth period [2, 3]. In the CBB cycle synthesized stromal 3-carbon phosphate esters (3-PGA (3-phosphoglycerate)) and triose phosphates (dihydroxyacetone phosphate, glyceraldehyde phosphate) are exported to the cytosol in exchange for P_i_ via the triose-P/P_i_ translocator or are utilized within chloroplasts for starch biosynthesis [4]. These changes not only affect chloroplast functions but also have broader implications as the balance between carbon assimilation, storage and utilisation is dependent on the partitioning of photoassimilates between starch and sucrose (suc) which itself is dependent on environmental conditions [4]. Given these metabolic complexities, it becomes essential to explore how regulatory mechanisms govern these processes. Subsequently, the regulation of carbon fixation and following carbohydrate metabolism is intricate and spans over various cellular compartments in plants [5, 6, 7, 8, 9]. Subsequently, regulatory mechanisms governing the processes between compartments involve transporters and their regulations within metabolic pathways [10, 11, 12, 13, 14]. Within the cytosol, a main metabolic route for the phosphorylated sugars is the synthesis of sucrose, one of the major transport forms of carbohydrates within plants. Sucrose is an important player of the energy metabolism and its synthesis is known to be a limiting factor during abiotic [15] and biotic [16] environmental perturbations.Sucrose itself is synthesized from hexose phosphates uridine diphosphate-glucose (UDP-glucose) and fructose-6-phosphate by sucrosephosphate synthase (SPS) and sucrose phosphate phosphatase (SPP, under P_i_ release) [17], while SPS marks the rate-limiting step of this reaction chain [18]. The regulation of SPS activity is multi-layered and involves phosphorylation that leads to inactivation [19] and is inhibited by UDP-glucose and P_i_. Activation is facilitated by e.g. glucose-6-phosphate (G6P), and to a lesser extent by glucose (glc) [20, 21, 22, 23, 24]. For example, the addition of G6P increased SPS activity 5-fold in *Oryza sativa* [25] and up to 16-fold in *Spinacia oleracea* [24]. Glucose activated SPS in *Ipomoea batatas* to a 1.4-fold increase [21]. For SPS inhibition, UDP-glucose is described for *Spinacia oleracea* to have a K_I_ of 9.4 [26]. Besides the interlaced regulation of SPS, sucrose can undergo the so-called sucrose cycling i.e. cyclic biosynthesis and degradation of sucrose which spans across various subcellular compartments like the cytosol, vacuole but also across the tonoplast [9, 15]. Either invertases catalyse the hydrolytic cleavage to glucose (glc) and fructose (frc), or, particularly in sink tissues, sucrose synthase (SuSy) cleaves sucrose into fructose and UDP-glucose or ADP-glucose [27]. Glucose and fructose can be rephosphorylated by hexokinases (HXKs, i.e. glucokinase and fructokinase), and the synthesized hexose phosphates serve again as substrate for sucrose biosynthesis (compare Fig. 1) [28, 29, 30]. Thereby, a continuous cycle is created, which introduces an additional regulatory strategy within the carbohydrate metabolism. Along with the regulation of SPS activity, the other enzymes within the cycle are also tightly regulated. Invertases are inhibited by glc and, to a lesser extent, by frc [31, 32]. For instance, cytosolic invertase (cINV) is inhibited by fructose and reaches only 20 % of its activity while glc led to only 10 % within a purified bamboo cell culture [32]. Additionally, both HXKs are subject to either inhibition by G6P or fructose-6-phosphate (F6P) [33]. In potato tubers HEXOKINASE1 (HXK1, a glucokinase (GlcK)) is non-competitively inhibited by G6P (K_I_ = 4 mM), while FRUCTOKINASE1 (FrcK) and FrcK2 are mostly non-competitively or partially competitive inhibited by F6P (K_I_ = 12 mM). These regulatory examples highlight the interwoven regulatory system of sucrose cycling. The cycling itself seems to be energetically wasteful (futile cycle), but it is considered to allow precise control over carbon partitioning [17], and estimations reach up to 10-30 % recycling flux within leaves and cotyledons for various species [34, 35, 36]. Additionally, it was shown via kinetic parameter estimations that HXKs seemed to be a regulator of this cycle, while invertase sucrose degradation was secondary [34, 37]. Supporting this, impairment of HXK1 in the *gin2-1* mutant indicates problems in the assimilate transport, shoot growth [38] and leads to sucrose accumulation and enhanced root respiration [39]. As a result, regulation of this interlaced cycle is of special interest because within or between compartments, cycles and their biochemical regulations help to maintain a metabolic balance during environmental perturbations [40, 9, 41, 42, 37]. In general, resolving cycles is still often challenging, as the stoichiometric information alone is frequently not enough to predict the dynamical behaviour of cellular systems [43, 44]. Therefore, such cycles are of special interest, but hard to elucidate, due to their complexity and a combination of computational biology and laboratory data is one way to shed light on this topic. To investigate sucrose cycling, we have chosen subcellular metabolite data in combination with mathematical modelling. We investigated system perturbations and stabilities by the structural kinetic modelling approach (SKM) [45]. With the SKM strategy, kinetic parameters are normalized to a steady-state condition enabling a fast simulation of millions of normalized enzyme kinetic parameter constellations. One advantage of SKM is, that only metabolic and network information are necessary to determine whether implemented regulations are stable within the pathway system or not [46, 45]. The method is frequently utilized for a range of research questions [45, 46, 47, 37, 48, 49] and assist in identifying key areas for further i.e. experimental investigations. A specific example is the predicted system instability in the sucrose cycling process caused by vacuolar regulation [37]. This prediction was confirmed experimentally, demonstrating that a deficiency in vacuolar invertase impacts photosystems during environmental stress conditions by sucrose cycling misregulation [9]. Since it has been suggested that sucrose cycling is regulated rather more by cytosolic HXKs than by cytosolic invertases [34, 37], we aimed to investigate this regulatory key lever by analyzing system stabilities in a subcellular model. We have chosen to investigate the HXK1 deficient mutant *gin2-1*. The mutant *gin2-1* (glucose insensitive) is impaired in its glucokinase activity (10 % of maximum activity from *Arabidopsis* Ler wild type [50, 51]) and is sensitive to high-light stress [50]. We investigated putative regulations and system stabilities in high-light and control conditions at two-time points based on available subcellular metabolite data [8, 51]. Our models specifically focused on differentiating between glc and frc and their potential distinct roles in sucrose cycling. We present mathematical modelling evidence that emphasizes the significant role of fructokinase activity and suggests a possible activation of SPS by fructose itself. As neither the BRENDA database [52] nor further investigated publications have documented SPS activation by fructose, we additionally confirmed this activation in vitro. Consequently, we propose that phosphorylated sugars as well as glc and frc serve as activators of SPS.

**Figure 1:**
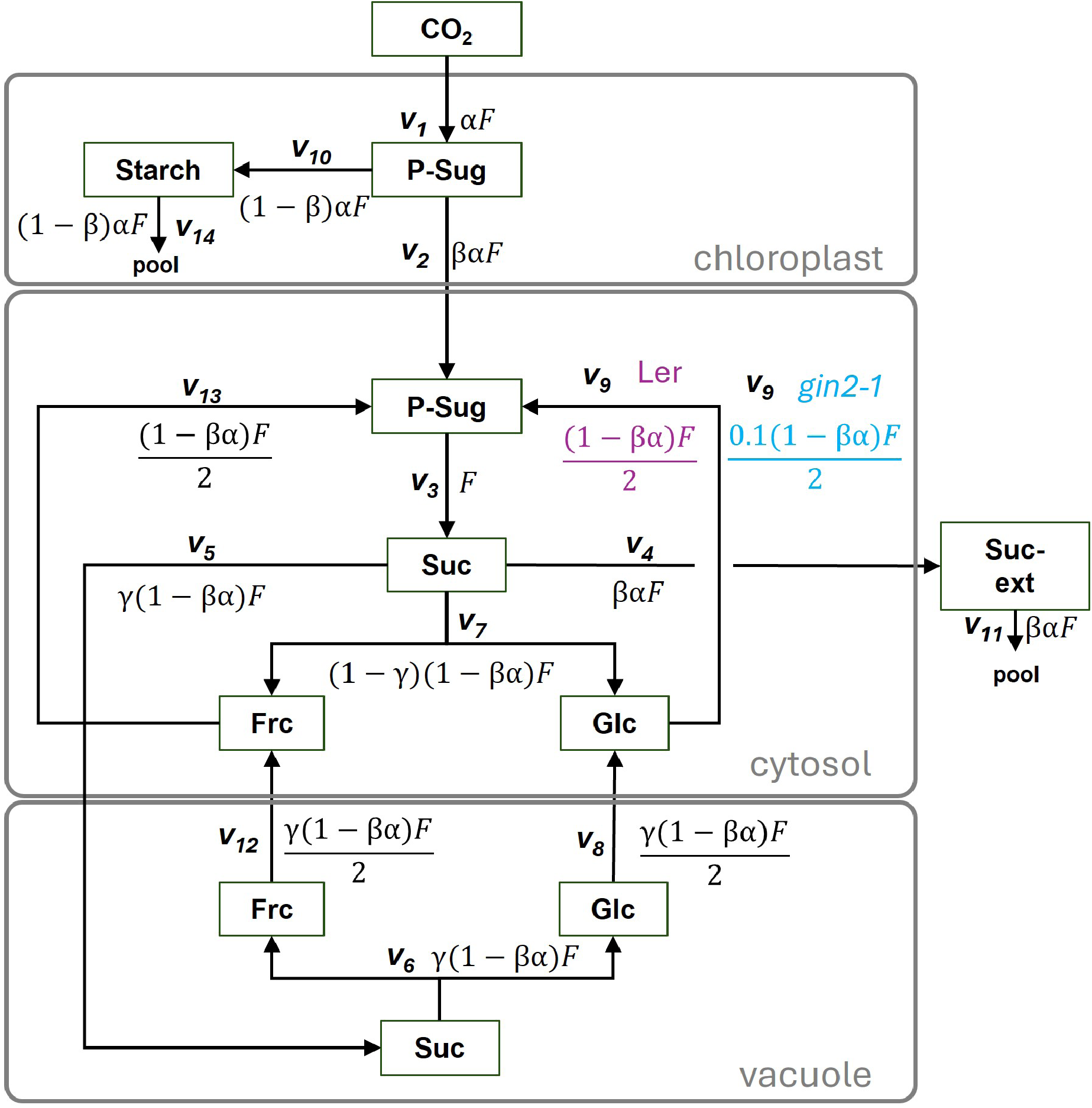
Structure of the leaf subcellular carbohydrate model. The model comprises the compartments of chloroplast, cytosol, vacuole and extracellular (ext) space. Metabolites in boxes are connected via reactions *v*_1_ to *v*_14_ and their corresponding fluxes. The following main-reactions are considered: *v*_1_:= *CO*_2_-fixation and resulting phosphorylated sugar pool; *v*_2_:= export of phosphorylated sugars; *v*_3_:= sucrose biosynthesis (sucrose-phosphate synthase, sucrose phosphate phosphatase); *v*_4_:= export of sucrose to extracellular space/other reactions; *v*_5_:= vacuolar sucrose import; *v*_6_/*v*_7_:= vacuolar/cytosolic invertase; *v*_8_/*v*_12_: vacuolar hexose export; *v*_9_:= glucokinase; for *gin2-1* 10%; *v*_10_:= starch biosynthesis; *v*_11_/*v*_14_:= pool of sucrose/ starch; *v*_13_:= fructokinase. Frc:= fructose, Glc:= glucose, P-Sug:= phosphorylated sugars, Suc:= sucrose, F:= flux, (*α*):= randomized perturbation factor, (*β, γ*):= randomized proportion factors for split-reactions.

## Material and Methods

### Model Preliminaries

To explore the reduction of glucokinase activity in *gin2-1* and the influence of individual hexoses (glc and frc) on the stability of the regulatory network (Fig. 1), we expanded a previous model [37].The model consists of a simplified carbon fixation process via phosphorylated sugar (P-Sug) formation, subsequent starch and sucrose biosynthesis and includes sucrose cycling (Fig. 1). Furthermore, we split vacuolar and cytosolic hexose pools into glc and frc, as well as separating hexokinases into glucokinase (GlcK) and fructokinase (FrcK). Additionally, we redefined the starch pool to function as sink during daylight hours. Fluxes were calculated; (*α*) was included as randomized perturbation factor, while (*β, γ*) served as randomized proportion factors for split-reactions. For *gin2-1* models we implemented a 10 % rest activity for the GlcK reaction (measurements from [38]). In total, the models comprised of 14 reactions (*v_j_*; *j* = 1,*…, r* here: *r* = 14), subcellular steady-state concentrations (*c*_0_*, i*; *i* = 1*, …, m* here: *m* = 10) and their corresponding fluxes *v_j_*(*c*_0*,i*_). The SKM approach [46, 45, 53] is based on analyzing the Eigenvalues of the Jacobian matrix (*J*), which itself is approximated by (equation 1):

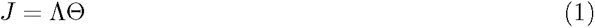

The Λ matrix (see Figure S1) is the stoichiometric matrix (*N*) normalized to steady state fluxes (*v*(*c*_0_)) and steady state metabolite concentrations *c*_0_ by the equation 2:

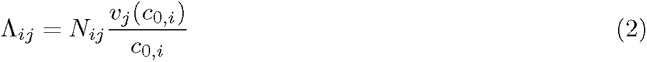

Entries of the matrix *θ* (equation 3, Figure S2) represent normalized elasticities, i.e. the degree of saturation of a normalized flux *µ* with regard to the normalized substrate concentration *x* (equation 3).

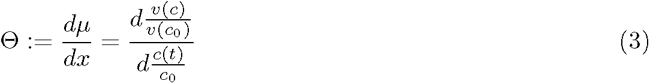

A non-zero entry in the Θ matrix indicates the involvement of a metabolite as a substrate (0; 1], as an activating (0; 1], or as an inhibiting compound [−1; 0). Finally, the Eigenvalues (*λ*s) of the Jacobian matrix *J* are calculated. The maximum real Eigenvalue is determined (*λ_max_* ∈ *R*) for each calculation. This *λ_max_*serves as a measure of stability. If *λ_max_ >* 0, then the system is defined as instable for this instance [46, 45]. In a biological context this can lead to extensive growth e.g. the metabolite pools reach non-physiological levels. If *λ_max_ <* 0, the system is defined as stable [46, 45] as extensive growth or decline of a metabolic pool results in positive real parts of Λ [49]. In our models (Fig. 1) the total flux *F* was set to 1 and was also tested for some models with higher flux rates (2 and 10 – examples are in Fig.S3). The distribution of splitting reactions was randomized by (*α, β,* and *γ*) within (0; 1). Similarly, for every calculation step, elasticities in Θ were randomized between (0; 1) except for those *θ*s of interest (see Fig. S2). This means every metabolite which is a substrate for a reaction also activates that reaction with a corresponding *θ*. This is especially important, as P-Sug activation of SPS is known from literature [24, 25] and also shown in our in vitro approach (Fig. 6). In this study, we defined the extent of instability by the proportion of positive maximal Eigenvalues across all iterations of every model. We tested 118 different combinations of “core regulations” for 10^6^ iterations. Models resulting in fewer than 100 instances (0.01%) of maximal positive Eigenvalues were termed “quasistable” to prevent over-interpretation of stabilities (false negative). For the strength of activations and inhibitions (entries in the Θ matrix) we used s=strong (100 % for activations and 99 % for inhibitions), m=medium (33 % for activations and 50 % for inhibitions), and w=weak (1 %). All models were calculated within the MATLAB software (The MathWorks, https://www.mathworks.com). The source code for the model is in GitLab (project ID 107389, https://git.rwth-aachen.de/Lisa.Fuertauer/skm-sucrose-cycling).

### Subcellular metabolite data input and initial testing

Subcellular metabolite data of glc, frc and suc were taken from *Arabidopsis thaliana* genotypes, *gin2-1* (HXK1 deficient mutant) and its corresponding background Landsberg erecta (L*er*) at different time points and conditions [8, 51]. Levels from starch and phosphorylated sugars (P-Sug, glucose-6-phosphate (G6P) and fructose-6-phosphate(F6P)) were taken from Küstner and colleagues [38]. Subcellular volumes were implemented by estimations of spinach leaves data [54] similar to previously [37]. The metabolic dataset is summarized in Supplementary Table 1. Roughly, the plants were grown for 6 weeks and harvested under control (120 *µ*mol m*^−^*^2^ s*^−^*^1^) and high-light (1200 *µ*mol m*^−^*^2^ s*^−^*^1^) conditions after 4-hour and 10-hour light periods. We set external suc concentrations to the same level as cytosolic suc, similar to a previous study [37]. Nevertheless, we tested external suc concentration (Fig. 1) to be either 2/3 (low), equal (medium), or 3/2 (high) compared to the internal cytosolic suc concentration. Notably, the external suc concentration did not majorly affect the number of instable instances for the tested models S3. Therefore, only datasets with medium external suc concentration were considered after initial analyses.

### Biochemical sucrose-phosphate synthase activity assay

For sucrose-phosphate synthase (SPS) activity determination, *Arabidopsis thaliana* Col-0 plants were cultivated for 5 weeks in a 12-hour day-night cycle, with day/night temperatures 22 °C / 18 °C. At midday, entire rosettes of five plants were harvested, pooled and frozen in liquid nitrogen and freeze-dried after grinding. The extraction and SPS assay were adapted from previously [55]. Shortly, freeze-dried plant material was incubated for 15 minutes on ice with an extraction buffer [50 mM HEPES-KOH (pH = 7.5), 10 mM MgCl_2_, 1 mM EDTA, 2.5 mM DTT, 10 % glycerol, and 0.1 % Triton-X-100]. Extracts were incubated with reaction buffer [50 mM HEPES-KOH (pH 7.5), 15 mM MgCl_2_, 2.5 mM DTT, 35 mM UDP-glucose and 35 mM fructose-6-phosphate (F6P)]. Additionally, to assess their ability to activate SPS reactions, 140 mM of either the putative activating sugars glucose-6-phosphate, glucose, fructose or pure water was added. The reaction was determined at 25 °C and terminated by addition of 30 % KOH and denaturing for 10 minutes at 95 °C. Blanks with direct termination were taken. Sucrose concentrations were measured via the anthrone assay [56, 57] and determined with a standard curve as well as measurements of a sample blank.

## Results

### *gin2-1* showed a higher extent of instability over all tested models

The expansion of the previous models [37, 47] allowed us to investigate the cytosolic part of sucrose cycling more in-depth. The stability of the model was tested without any further regulations and resulted, as expected, in solely negative *λ_max_* values (Fig. S4). This allowed us to investigate different regulation scenarios. We initially tested strategically different activation and inhibition combinations, and followed with a combinatorial manner based on results from previous model outputs. In summary, we tested 118 models with different activation and inhibition scenarios for all genotypes, conditions and timepoints (Fig. 2 A). These regulations consisted of inhibitions of glucokinase (GlcK)/fructokinase (FrcK) by phosphorylated sugars (P-Sug), the activation of SPS by hexoses and P-Sug, the inhibition of cytosolic and vacuolar invertases by the hexoses, and the activation of sucrose import into the vacuole. Additionally, we tested for weak (*θ* = –0.01), medium (*θ* = –0.5), and strong (*θ* = –0.99) inhibitions, as well as strong (*θ* = 1) and medium (*θ* = 0.33) activations within the cytosol as well as the vacuole (Fig. 2 A). In the majority of the models, similar patterns of stability/instability were found in both genotypes (Fig. 2 B, green and red). Approximately half of our models were instable for both genotypes (Fig. 2 B, red). Analysis of differentiating cases revealed, that all regulations, which led to instability for Ler, but stability for *gin2-1* in one condition (Fig. 2 B, turquoise) also showed stability for Ler but instability for *gin2-1* in another condition (Fig. 2 B, Supplementary Table 2). In total, 13 differentiating models were discovered, which dissolve into either stabilization or destabilization from one condition to another (Fig. 2 B). Furthermore, we determined the total number of instable cases (*N_λ_ >* 0) within the 10^6^ runs and compared them between the two genotypes. In over 80 % of cases for the control (C) and all cases for high-light (HL), *N* was higher in *gin2-1* (Fig. 2 C). Additionally, for the control cases, the proportion increased for the 10 h time point. To elucidate the mechanistic difference between *gin2-1* and wild type stability (Fig. 2 B/C, turquoise, purple), the 13 differentiating models were examined for common characteristics: 12 of these models were combinations of multiple SPS activators, either by double glc and frc activation (6 regulations) or triple glc, frc and P-Sug activation (6 regulations) (see Supplementary Table 2).

**Figure 2:**
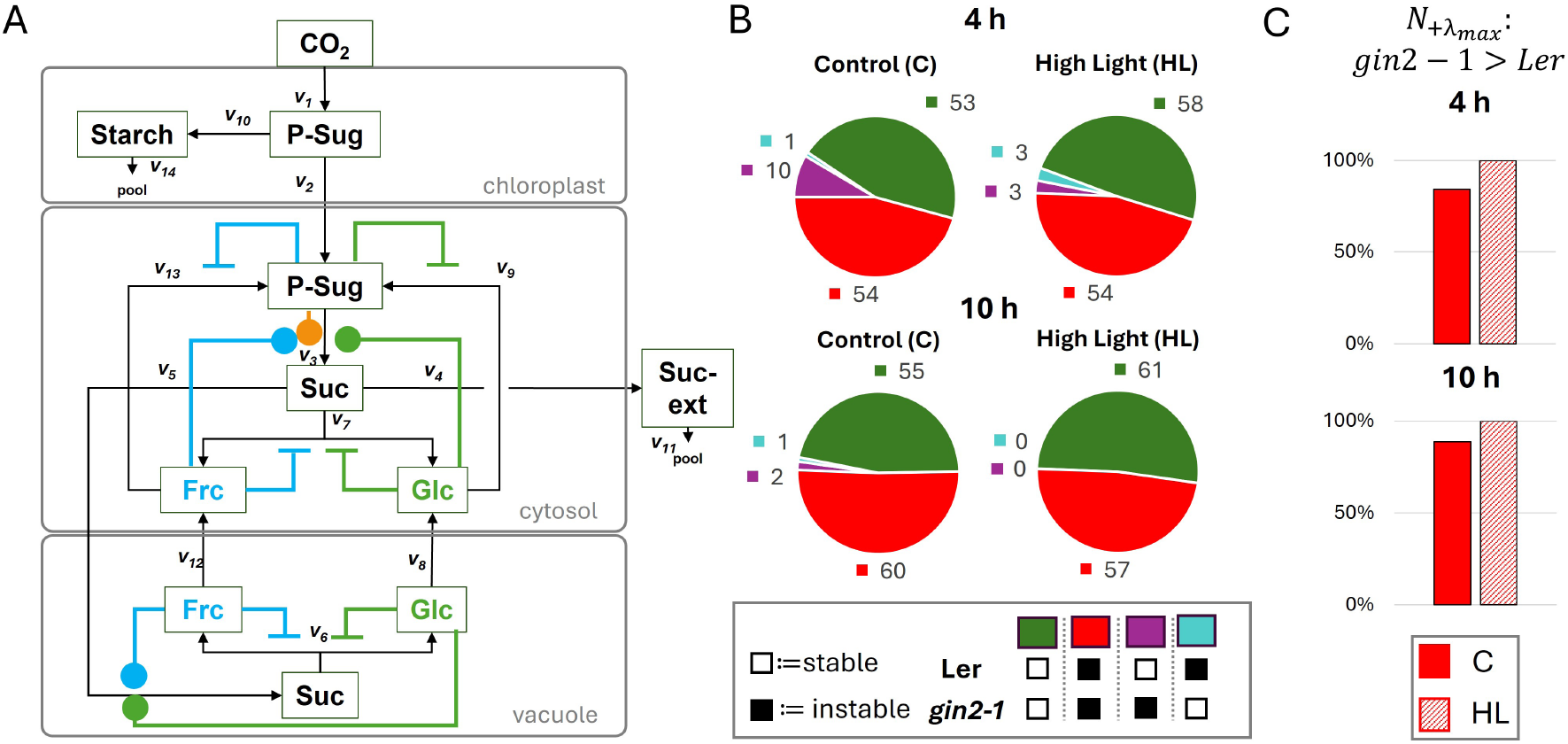
Overview of all implemented regulations and summary of all results. A) 118 models with different combinations of activations (blue/green/orange circles) and inhibitions (blue/green bars) were tested. A detailed summary with all combinations and results can be found in Supplementary Table 2. B) Overview of stability status of SKM model runs for 4 h, 10 h, control, and high-light conditions in both genotypes. Color code indicates stability (white box) and instability (black box) in genotypes. Models with stability for Ler are in green and purple, for *gin2-1* in green and turquoise. C) The instable models for both genotypes (B, red) were compared between the two genotypes. The individual models were analysed in 10^6^ runs. The total number *N* of positive +*λ_max_*was determined and compared between the two genotypes. In the majority of cases for the control (C), and in all cases for high-light (HL), *N* was higher in *gin2-1*.

### Fructokinase inhibition leads to higher *gin2-1* instability

The incorporation of double activation by glc and frc on SPS led to a stable model for both genotypes and all conditions, if both hexokinases (by P-Sug) and cytosolic invertases (by glc+frc) were strongly inhibited (Fig. 3 A). For single hexokinase inhibition scenarios, instabilities arise (Fig. 3 B-G). In detail, the GlcK inhibition cases led to instable solutions for both genotypes and conditions (Fig. 3 B-D), whereas the FrcK inhibition cases led to higher stabilities, especially in Ler (Fig. 3 E-G).

**Figure 3:**
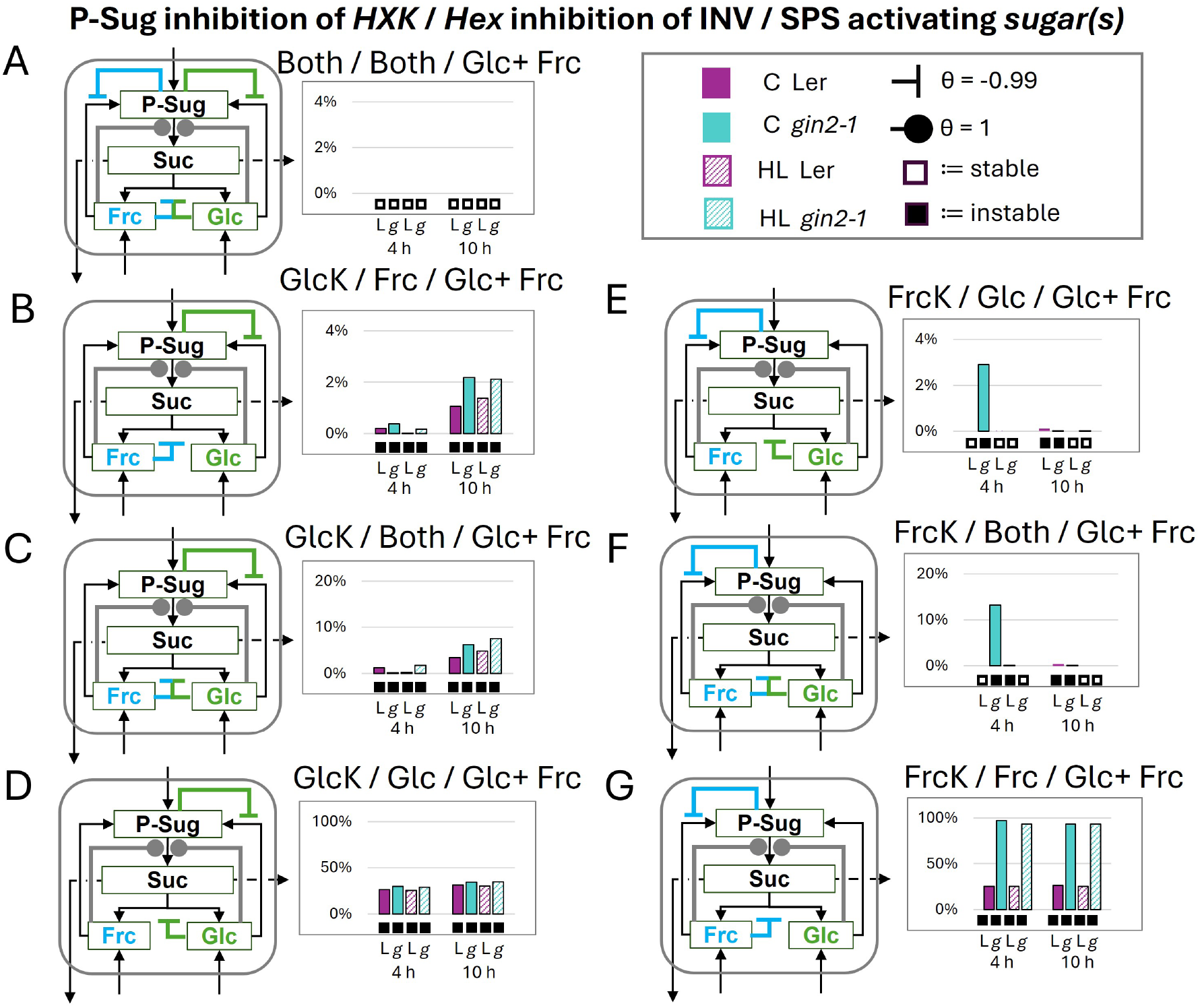
Influence of invertase and hexokinase inhibitions on the cytosolic sucrose cycle stability by double glucose and fructose SPS activation. Stability analysis of full SPS activation by both non-phosphorylated hexoses. The subsequent combinations are indicated above each diagram as follows: inhibition of P-Sug on which *hexokinase* / which *hexose* inhibits cytosolic invertase / which *sugar(s)* activates SPS. A) all inhibitions and activation scenarios (B–D) glucokinase inhibition (E–G) fructokinase inhibition with varied invertase inhibition scenarios. Filled circles indicate activation and inhibition is indicated by inhibition arcs, specific *θ*s are given in the figure. Boxes below diagrams indicate stability (empty box) and instability (black box). Note: Here only the cytosolic regulations are depicted, but the model consisted of the whole cell (Fig. 1). L:= Ler (purple, control/high-light:= filled/crosshatched), g:= *gin2-1* (turquoise, control (C)/high-light (HL):= filled/crosshatched), HXK:= hexokinase (GlcK:=glucokinase, FrcK:=fructokinase), INV:= invertase (here cytosolic), Frc:= fructose, Glc:= glucose, P-Sug:= phosphorylated sugars, SPS:= sucrose-phosphate-synthase, Suc:= sucrose

Additionally, FrcK inhibition seemed to be more stable in 10 h high-light conditions (Fig. 3 E/F). Interestingly, if the implemented HXK inhibition did not correspond with the hexose, which inhibited the cytosolic invertase (Fig. 3 B/E), the extent of instability only slightly increased to maximal 3 %. If additionally the second hexose inhibited the invertase, the extent of instability increased (Fig. 3 C/F). Here, the 10 h time point showed higher extents between 3-7 % for the glucokinase inhibition cases (Fig. 3 C). For fructokinase inhibition, the highest extent of instability with 13 % was reached by *gin2-1* at 4 h under control conditions (Fig. 3 F). In the case of Ler also the 4 h high-light condition was then instable (Fig. 3 E vs F). Instability increased for all genotypes and conditions, if the inhibited hexokinase corresponded to the invertase inhibiting hexose (Fig. 3 D/G). Here, all genotypes and conditions reached at least 20 % for their extent of instability. For *gin2-1* instability increased to at least 90 % for both time points and conditions in the fructokinase inhibited case (Fig. 3 G), while it was unchanged compared to glucokinase inhibition in Ler (Fig. 3 D/G). All preceding effects could be dampened by adding an additional weak inhibition of the not-yet inhibited hexokinase, but still the stability patterns stayed the same (Fig. S5).

### Glucose and fructose as potential activators of SPS

Since the SPS double activation by glc and frc without double feedback inhibitions of INV and HXK led to instability (Fig. 3 B-G), we were interested in the effect of a single activation of SPS by glc or frc (Fig. 4). If the reaction network was regulated one-sided within the model (Fig. 4 A/E), it was stable for both glc and frc SPS activation. This is not the case with double activation (compare Fig. 3 D/G). Inhibiting the invertase with the non-activating hexose i.e. the GlcK/Frc or FrcK/Glc system (P-Sug inhibition of HXK/Hex inhibition of INV) (Fig. 4 B/F) resulted in a slight increase in instability. In contrast, instability was found when the implemented regulation was contrary within the cycle, i.e. the hexose, which inhibits the invertase and activates SPS, does not correspond to its inhibited HXK (Fig. 4 C/G). Again, *gin2-1* showed higher instabilities (30 to 40 %) compared to Ler (20 %). The highest instabilities were found when both inhibitions did not correspond with the SPS activator (Fig. 4 D/H). Here, instability increased to around 60 % for the wild type in both activator cases. For *gin2-1* extents of around 70 % in the frc activator case (Fig 4 D) and total instability (100 %) was determined for the glc activator case (Fig 4 H). Since the wild type behaved similarly for both activator cases, frc and glc show the same ability to activate the SPS according to SKM. As P-Sug are a substrate for the SPS reaction, the theta entry within the matrix-system is always positive and therefore it can be interpreted as activating the subsequent reaction. Testing for a strong activation of SPS by P-Sug (*θ* = 1) led to only stable solutions for previously described combinations (P-Sug inhibition on HXKs and feedback inhibition on INV from glc and/or frc) (Fig. S6 A-D).

**Figure 4:**
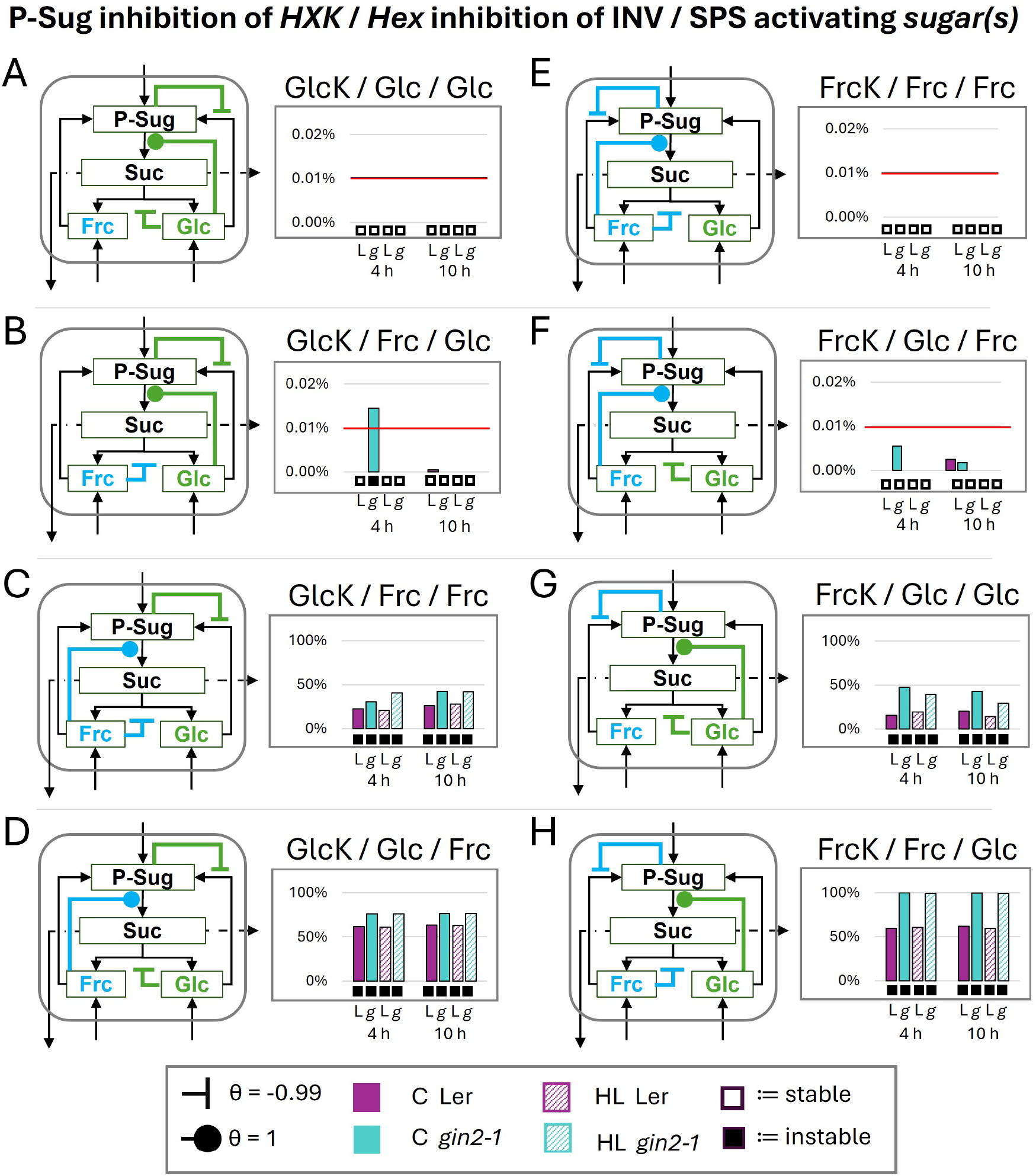
Glucose or fructose as putative activators of SPS. Stability analysis of full SPS activation by single hexoses. The subsequent combinations are indicated above each diagram as follows: inhibition of P-Sug on which *hexokinase* / which *hexose* inhibits cytosolic invertase / which *sugar(s)* activates SPS. (A-D) Glucokinase inhibition with (A-B) glucose and (C-D) fructose as SPS activators. (E-H) Fructokinase inhibition with (E-F) fructose and (G-H) glucose as SPS activators. Filled circles indicate activation and inhibition is indicated by inhibition arcs, specific *θ*s are given in the figure. Boxes below diagrams indicate stability (empty box) and instability (black box). Note: Here only the cytosolic regulations are depicted, but the model consisted of the whole cell (Fig. 1). L:= Ler (purple, control/high-light:= filled/crosshatched), g:= *gin2-1* (turquoise, control (C)/high-light (HL):= filled/crosshatched), HXK:= hexokinase (GlcK:=glucokinase, FrcK:=fructokinase), INV:= invertase (here cytosolic), Frc:= fructose, Glc:= glucose, P-Sug:= phosphorylated sugars, Suc:= sucrose, SPS:= sucrose-phosphate-synthase; Red line cut-off for instability

### Triple activation of SPS can lead to fewer instabilities

In our next setup we tested the triple activation of SPS by glc, frc and P-Sug. Our previous observations have shown that on the one hand both non-phosphorylating hexoses were able to activate the SPS (Fig. 4) and on the other hand in our setup P-Sug as SPS activator only lead to stable solutions (Fig. S6). In contrast to the prior investigations also weak HXK inhibitions were added, since these showed also a stabilizing effect for the double activation by both non-phosphorylated hexoses (Fig. S5). Also, the activation power of the non-phosphorylated hexoses was set to a lower level (*θ* = 0.33), since P-Sug was more robust as SPS activator according to our SKM data and also G6P is stronger in its activation capacity than glucose according to in vitro data.

Overall, the extents of instabilities were lowered in all tested combinations in comparison to the SPS activation by the single (Fig. 4) and double (Fig. 3) activation of the non-phosphorylated hexoses, even with the additional weak HXK inhibition (Fig. S5). Still, the same pattern of increasing instability extent going from non-corresponding inhibitions to corresponding inhibitions in respect of the non-phosphorylated hexoses could be observed (Fig. 5 D/G). Moreover, the higher impact of fructokinase inhibition on *gin2-1* stability in comparison to wild type was also detected (Fig. 5 D/G).

**Figure 5:**
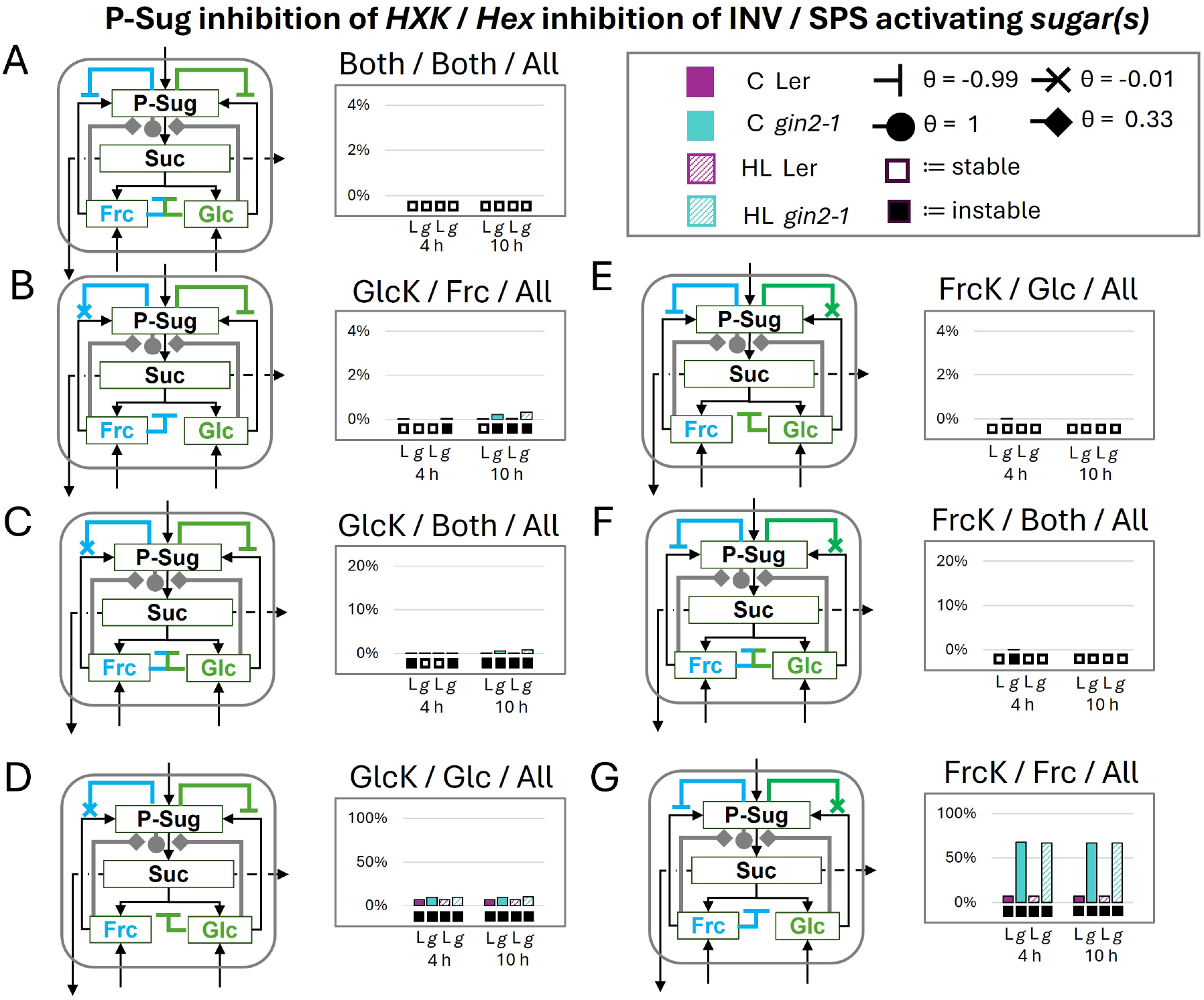
Influence of triple activated SPS. Stability analysis of full SPS activation by phosphorylated sugars (P-Sug) and minor activation by the non-phosphorylated hexoses. The subsequent combinations are indicated above each diagram as follows: strong inhibition of P-Sug on which *hexokinase* / which *hexose* inhibits cytosolic invertase / which *sugar(s)* activates SPS. The regulation combinations here are based on the regulations including weak hexokinase inhibitions (Fig. S5). A) all inhibitions and activation scenarios (B – D) strong glucokinase and weak fructokinase inhibition (E – G) strong fructokinase and weak glucokinase inhibition. Filled circles and diamonds indicate activation and inhibition is indicated by inhibition arcs and crosses, specific *θ*s are given in the figure. Boxes below diagrams indicate stability (empty box) and instability (black box). Note: Here only the cytosolic regulations are depicted, but the model consisted of the whole cell (Fig. 1). L:= Ler (purple, control/high-light:= filled/crosshatched), g:= *gin2-1* (turquoise, control (C)/high-light (HL):= filled/crosshatched), HXK:= hexokinase (GlcK:=glucokinase, FrcK:=fructokinase), INV:= invertase (here cytosolic), Frc:= fructose, Glc:= glucose, P-Sug:= phosphorylated sugars, SPS:= sucrose-phosphate-synthase, Suc:= sucrose

### SPS can be activated by fructose according to in vitro ***v_max_*** activity data

Since the triple activation of SPS resulted in the lowest extents of instability, but still showed a difference between Ler and *gin2-1*, this prediction was tested in vitro: For P-Sug, such as glucose-6-phosphate (G6P), strong activation [24, 25] and for glc mild activation [21] of plant-derived SPS were previously demonstrated in vitro. However, the activation capability of frc, to our knowledge, and according to the BRENDA database [52] is barely considered. Given the in silico suggestion that frc could also potentially activate SPS, an in vitro SPS activity assay was conducted using leaf rosette extract of Col-0 *Arabidopsis thaliana*. We either added G6P, glc, or frc as the activating compound and used water as a control, to determine the effects of these components on the plant’s SPS (Fig. 6). As expected, G6P emerged as the strongest activator, leading to an activity of around 54 µmol suc/(gDM*h), while glucose only led to an activity of around 19 µmol suc/(gDM*h). Addition of frc led to a similar activity as glc with ca. 23 µmol suc/(gDM*h). All three seem to exhibit activation capabilities, since the water control was at a lower activity level with around 5 µmol suc/(gDM*h).

**Figure 6:**
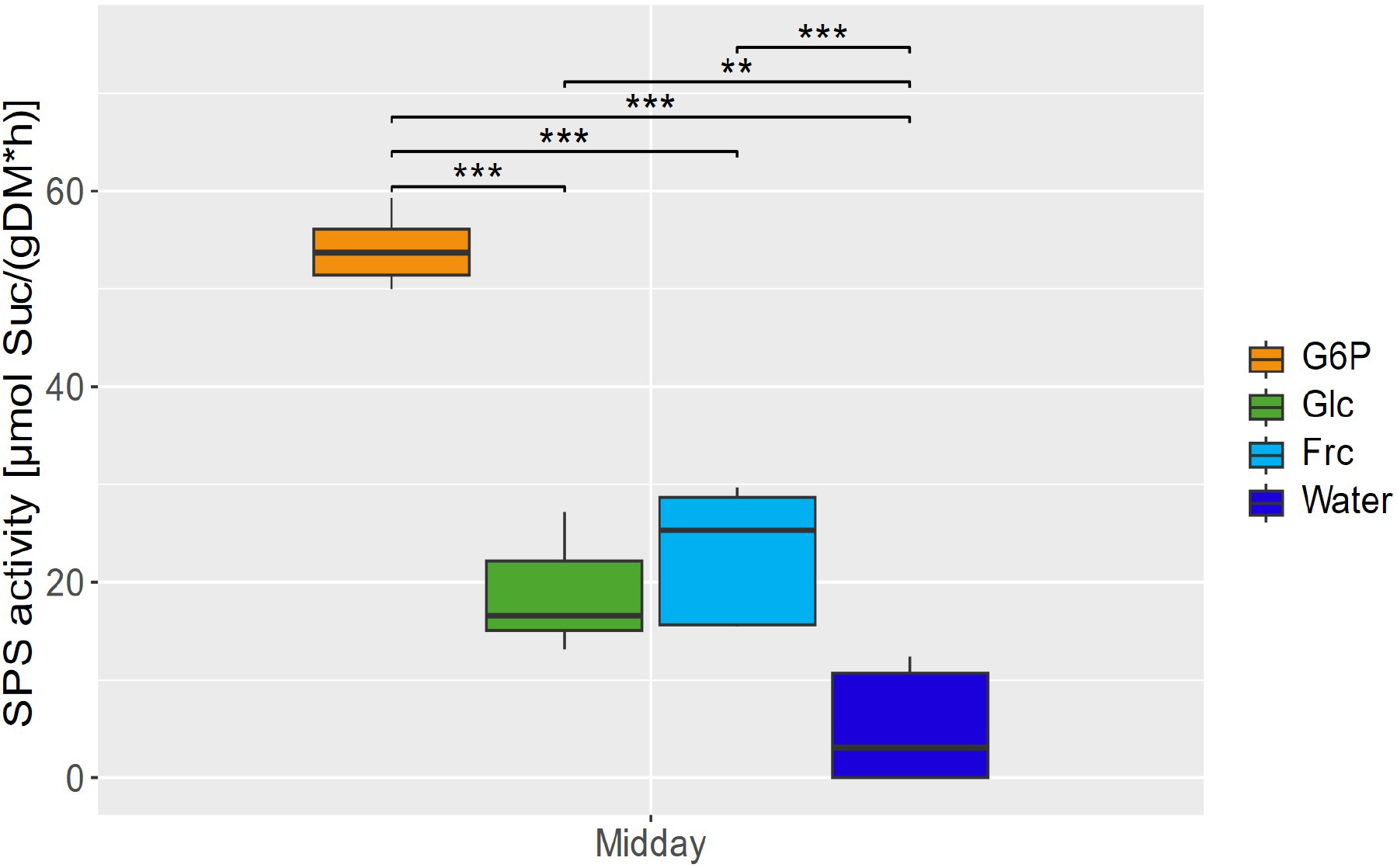
In vitro determination of SPS activity with activation by sugars in crude *Arabidopsis* Col-0 extracts. The sugars glucose-6-phosphate (G6P), glucose (Glc) and fructose (Frc) were independently added to crude leaf rosette extracts and the maximum SPS activity was deter-mined. For comparison, a water control without additional sugars was determined. **p *<* 0.01, ***p *<* 0.001 (t-test).

## Discussion

### High-light stress sensitivity of *gin2-1* is mathematically confirmed by SKM

HEXOKINASE1 is known to play an intrinsic part of the plant glucose signaling network [50]. Yet, the HXK1 deficient mutant *gin2-1* lacks catalytic and sugar sensing/signalling function of glucokinase and shows growth impairment together with being sensitive to high-light conditions [50, 51]. SKM stability analysis allowed us to test for 118 different combinations within the sucrose cycling (Fig. 2 B) and the two genotypes, two time points and conditions. Interestingly, in some cases, high-light conditions seemed to stabilize some of the investigated systems (Fig. 2 B). An explanation for this effect might be the accumulation of soluble sugars under high-light stress (Supplementary Table 1). Here, within the SKM system, it is similar to Michaelis-Menten enzyme kinetics. A higher concentration of the activator and/or inhibitor can drastically influence the enzymatic reaction rates *v_max_* as well as the *K_m_*. We observed in the majority of the models similar patterns of stability/instability for both genotypes (Fig. 2 B). Regarding the absence of a clear divergence of instability patterns between the genotypes, one possible explanation is that *gin2-1* mutants can still thrive under the tested conditions and that the mutant is not embryo-lethal. Nevertheless, what we could observe were the higher extents of instability in *gin2-1* compared to the wild type (Fig. 2 C). For high-light, this difference was fully pronounced since under this condition all instable models resulted in higher extents of instability for the mutant (Fig. 2 C). We interpret this as a sign, that with 10 % rest-activity of GlcK, the systems are more challenged under high-light conditions and confirm mathematically the experimentally observed high-light sensitivity of *gin2-1*. Those challenges could be, besides of a higher need for signalling function of HXK1, also an increased sucrose cycling activity. As discussed above, increased sucrose cycling activity might be a way to allow precise control over carbon partitioning [17]. It seems to play an essential role under environmental challenges [37, 9, 34, 35, 36], and ought to be a system for energy balancing [15]. Our results also nicely correlates with the fact, that *gin2-1* seems to metabolically react slower in light conditions compared to its wild type [8, 51]. Additionally, inhibition of FrcK in *gin2-1* models, often led to higher instability levels, indicating a dependency on this alternative route. This could be another explanation for the lower stress tolerance of this genotype.

### Glucose and fructose as activators of SPS

Double activation of SPS by glc and frc, and therefore enhanced sucrose synthesis flux, led to system stability if strong double inhibitions on INV and HXKs were present (Fig. 3 A). Thus, strong inhibitions seemed necessary, as otherwise, the systems showed increased instabilities (Fig. 3 B-G). Here, the dependency of the sucrose cycling is shown for glucokinase, as all combinations were instable when glucokinase was inhibited by P-Sug (Fig. 3 B-D). This confirms previous observations, as glucokinase seems to play a major role within the sucrose cycling system [47, 34, 39]. In contrast, inhibiting fructokinase by P-Sug seemed to be problematic only for several cases and especially in *gin2-1* (Fig. 3 E-G). Interestingly, the mutant showed higher extents of instability at the 4 h control time points. Both genotypes revealed instabilities when fructokinase was inhibited and fructose acted as a strong invertase inhibitor (Fig. 3 G). These instabilities were nearly up to 100 % in *gin2-1*, which could indicate a high dependence on fructokinase as a regulator by lacking and/or downregulated fluxes through the glucose side. Since the double high activation of glc and frc on SPS led to instabilities, we wanted to check if the system is stabilized by single P-Sug (Fig. S6), glc or frc (Fig. 4) activation. Single activation of P-Sug always led to stable results (Fig. S6) which was also expected as P-Sug is a substrate for SPS reactions. For glc and frc, one-sided regulation led to stability (Fig. 4 A/E), which we interpret as a reflection of a coordinated and guidance of fluxes. Further analysis revealed, that the systems tended to be stabilized, if the activating hexose did have in parallel a strong inhibition on its corresponding HXKs (Fig. 4 B/F). In comparison, diverging the activating hexose from its HXK led to instability and the wild type behaved similarly for both activator cases (Fig. 4 C-H), and it seemed that single frc and glc show similar abilities to activate the SPS according to SKM.

### Triple activation of SPS – a regulation strategy for the sucrose cycle

Activation of SPS by P-Sug led to system stability (Fig. S6). On the other hand we observed a conserved stable regulation for double and single activation of SPS with parallel feedback on HXKs and cytosolic INV (Fig. 3 A, Fig. 4 A/E Supplementary Table 2). As multiple strong activations normally perturb the stability of the systems, we incorporated in the triple activation model a higher activation potential for P-Sug than for glc and frc (Fig. 5) as we estimated a higher activation potential from P-Sug on SPS than from hexoses from biochemical data. Indeed, these additions led to lower extents of instability, but the model kept its distinguishing nature and stability pattern for activation of glc or frc and the feedback on the corresponding HXK (Fig. 5 D/G). We concluded that this model – all three SPS activating sugars in combination, both HXKs strongly inhibited by P-Sug and both hexoses strongly inhibiting the cytosolic INV (Fig. 5 A) – could be close to the regulation within the phototrophic plant cells in vivo. Since SPS activation has been only shown in vitro for G6P [25, 24] and glc [21] according to the BRENDA database [52], we tested frc for its SPS activation capability (Fig 6). We were able to confirm that indeed frc is as capable as glc to activate SPS activity in vitro from crude extracts. For the determination of specific kinetic parameters, purification of the enzyme would be necessary. Nevertheless, we can support the data from [20, 21] for glc being an SPS activator and adding here frc as well. Additionally, the already described higher activation power of G6P could be determined [25, 24, 21], which also fitted with the more flexible SPS activation by P-Sug according to the stability results. Interestingly, a recent study concluded that SPS derived from *A. thaliana* leaves were not allosterically activated by G6P [58]. Since this study was performed with recombinantly expressed SPS in yeast, missing post-translational modifications like the phosphorylation could explain the lack of G6P activation for both shoot allocated SPS isoforms SPSA1 and SPSC. Nevertheless, the previously measured G6P mediated activation of leaf material SPS in two other organisms [25, 24] is a strong indicator that this is also the case in *Arabidopsis*. G6P potentially activates SPS to a greater level (approx. 3-fold) than glc and frc (Fig. 6). The weaker activation might be explained by the conditions required for a stable system. The comparison between the three activators (G6P, glc, and frc) showed a dependency of the non-phosphorylated hexose as an activator on corresponding HXK and INV inhibition (Fig. 4). Conversely, the HXK inhibition and the invertase inhibition must contradict, if both hexoses activate SPS (3). This complements our observation regarding single activator results (Fig. 4) since one corresponding inhibition per activator is needed to stabilize the system. Furthermore, these different stable states for single and double activation could hint to a self-balancing system when combined. Nonetheless, both types of inhibition do not contribute to the stability of the activator to the same extent: the inhibition of the HXK seems to have a stronger influence than the inhibition of the INV. This could be due to the shared fluxes for the invertase and two different fluxes for the HXKs in our system. For future investigation, it would be interesting to see if the implementation of sucrose synthase (SuSy) supports this hypothesis. Another possibility is, that the HXK reaction products, phosphorylated sugars, are activated sugars and therefore more tightly regulated. Additionally, none of the implemented biochemical regulations (SPS activations, HXK inhibitions and INV inhibitions) were taken from *Arabidopsis thaliana* data, but from several plant species [25, 24, 21, 26, 32, 33]. Since the central carbohydrate metabolism is well conserved, our SKM derived prognoses can be expected to be conserved over a wide span of plant species.

### Fructose as an additional regulator of SPS activity

Besides our SKM results, we interpret the biochemical results that frc stabilizes sucrose cycling. The shown capability of frc to activate SPS might be necessary for a robust sucrose cycle to avoid unwanted accumulations or reductions of both hexose pools, because phosphorylated sugars can inhibit both HXKs and both hexoses can inhibit invertases. The previously described interdependence of these regulations could explain the need for additional SPS activation by frc. This might be the reason why the combination of all three activators showed the lowest extents of instability (Fig. 5). Fructose could support or act as a signal for phosphorylated sugars such as fructose 2,6-bisphosphate, which is considered a controlling unit for partitioning of photoassimaltes and sucrose synthesis [4]. Moreover, SuSy can split suc into frc and UDP-glc or ADP-glc instead of glc [59]. If SuSy is more active, e.g. in sink organs, a higher frc pool compared to glc pool could arise, thus intrinsic frc signalling pathways are needed. Further, suc signalling in plants and suc export [60] could be supported by frc. It can also be speculated, that frc has a control function as an activator, if either FrcK or GlcK is disturbed (Fig. 4). Additionally, it was shown that when cytosolic frc levels are modified by transporters, plant organ development and stress tolerance are severely impacted [61]. Considering this, an expanded model, that includes the lower extent phosphorylation of fructose by HXK1 [62], could provide more insight into the complexity of frc as a regulator. In future investigations involving different mutations and combination of mutants within the sucrose cycle —ideally those affecting the allosteric centers of SPS or HXKs or INVs— could provide deeper insights into regulation of the sucrose cycle. In a characterization study, the putative allosteric center for G6P was located in the N-terminus. A truncation led to higher activity and might also hint at a self-inhibitory function of this domain [63]. A metabolic dataset derived by a mutant lacking this N-terminus could yield more insights and help for validation. Yet, this proposition regarding the allosteric region needs further validation. These further experiments could help to substantiate our proposed model. However, frc supports the non-intuitive signalling within plants. This study highlights again the strength of a systems biology approach by combining mathematical modelling techniques with biochemical analysis.

## Conclusions

In this study, we constructed a subcellular model of the plant carbohydrate metabolism with the goal of understanding the regulation of the sucrose cycling. We have identified differences in systems stabilities in the HEXOKINASE1 deficient mutant *gin2-1*. Additionally, we predicted mathematically that fructose can activate sucrose-phosphate synthase. This suggested that fructose is a core element in regulating the sucrose cycle. Subsequently, we were able to confirm fructose to be able to activate SPS in vitro. With that, we propose the biochemical regulation of this cycle including a triple activation by P-Sug, glc and frc. In summary, our results further strengthen the theory that cytosolic fructose is an underestimated player in sugar signalling networks.

Not applicable

## Availability of data and materials

The model code as well as calculated Eigenvalues can be found in the GitLab repository (project ID 107389, https://git.rwth-aachen.de/Lisa.Fuertauer/skm-sucrose-cycling). All further material is provided in the supplements.

## Competing interests

The authors declare that they have no competing interests.

## Funding

This project was funded by the WISNA program.

## Author contributions

LF conceived the study. LF and OG constructed the models. OG performed modelling and CB performed the biochemical assays. All authors contributed to writing the manuscript.

## Acknowledgements

We thank Angelina Jordine, Martin Siedt, Joost van Dongen, Matthias Freh-Jordine, and the members of the Fürtauer, van Dongen and Wormit groups for helpful discussions. This project was funded by the WISNA program.

## List of abbreviations

3-PGA: 3-phosphoglycerate
ADP-glucose: adenosine diphosphate glucose
CBB: Calvin-Benson-Bassham (cycle)
cINV: cytosolic invertase
DM: dry matter
UDP-glucose: uridine diphosphate glucose
F6P: fructose-6-phosphate
frc: fructose
FrcK: fructokinase
FrcK1: fructokinase 1
FrcK2: fructokinase 2
G6P: glucose-6-phosphate
*gin2-1*: glucose insensitive 2-1
glc: glucose
GlcK: glucokinase
HXK: hexokinase
HXK1: HEXOKINASE1
INV: invertase
Ler: Landsberg *erecta*
P-Sug: phosphorylated sugars
SKM: structural kinetic modeling
SPP: sucrose phosphate phosphatase
SPS: sucrose-phosphate synthase
SPSA1: sucrose-phosphate synthase A1 isoform
SPSC: sucrose-phosphate synthase C isoform
suc: sucrose
SuSy: sucrose synthase

## Supporting Information

Fig. S1 Λ matrix of the model.

Fig. S2 Θ matrix of the model.

Fig. S3 Stability analysis overview of different F flux sizes and external sucrose concentrations.

Fig. S4 Histograms of Eigenvalue real part maxima.

Fig. S5 Influence of additional weak hexokinase inhibitions to prior regulations on the cytosolic sucrose cycle stability.

Fig. S6 Influence of single activated sucrose-phosphate synthase (SPS) by phosphorylated sugars on the cytosolic sucrose cycle stability.

Supp. Table 1 Calculated subcellular concentrations of metabolites in Ler and *gin2-1* for all here tested conditions

Supp. Table 2 Proportions of positive maximal Eigenvalues for all tested regulations

https://git.rwth-aachen.de/Lisa.Fuertauer/skm-sucrose-cycling. GitLab Project ID 107389

## A Supplementary information

**Figure S1:**
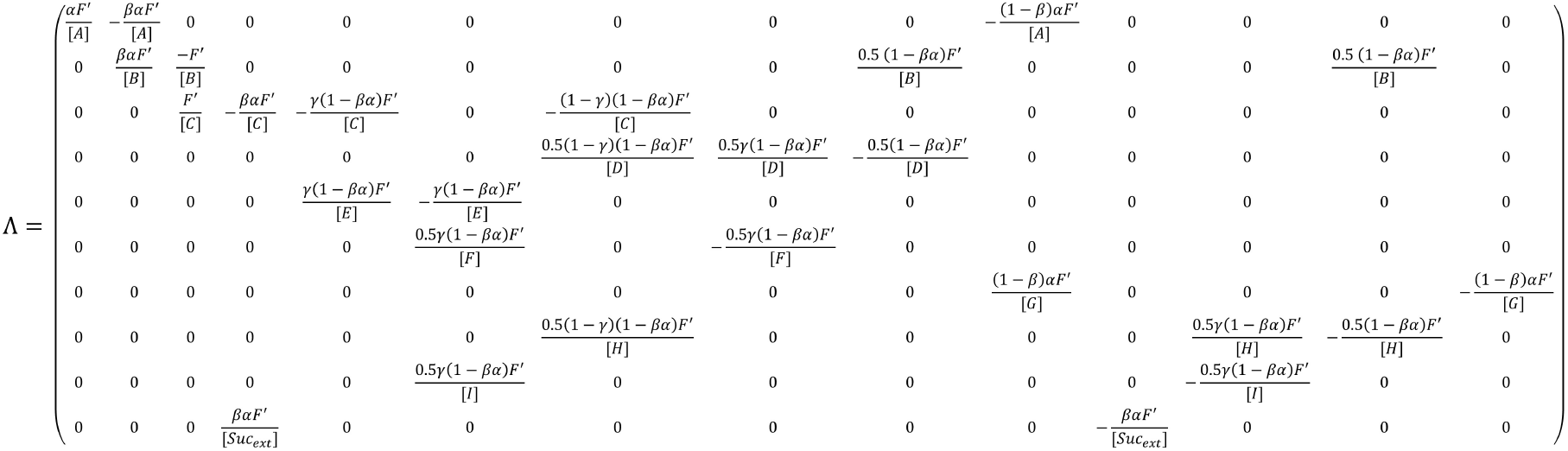
Λ matrix of the model. The matrix consists of 10 metabolite pools (rows): A: = P-Sugplastidic, B: = P-Sugcytosolic, C: = Succytosolic, D: = Glccytosolic, E: = Sucvacuolar, F: = Glcvacuolar, G: = starch_plastidic_, H: = Frc_cytosolic_, I: = Frc_vacuolar_ and 14 fluxes (columns). The concentrations depicted in the denominator are taken from the metabolic dataset. (*α*), (*β*) and (*γ*) were randomized between 0 and 1 for every simulation run. F was set to 1, but also F = 2 and F = 10 were initially tested (Fig S4).

**Figure S2:**
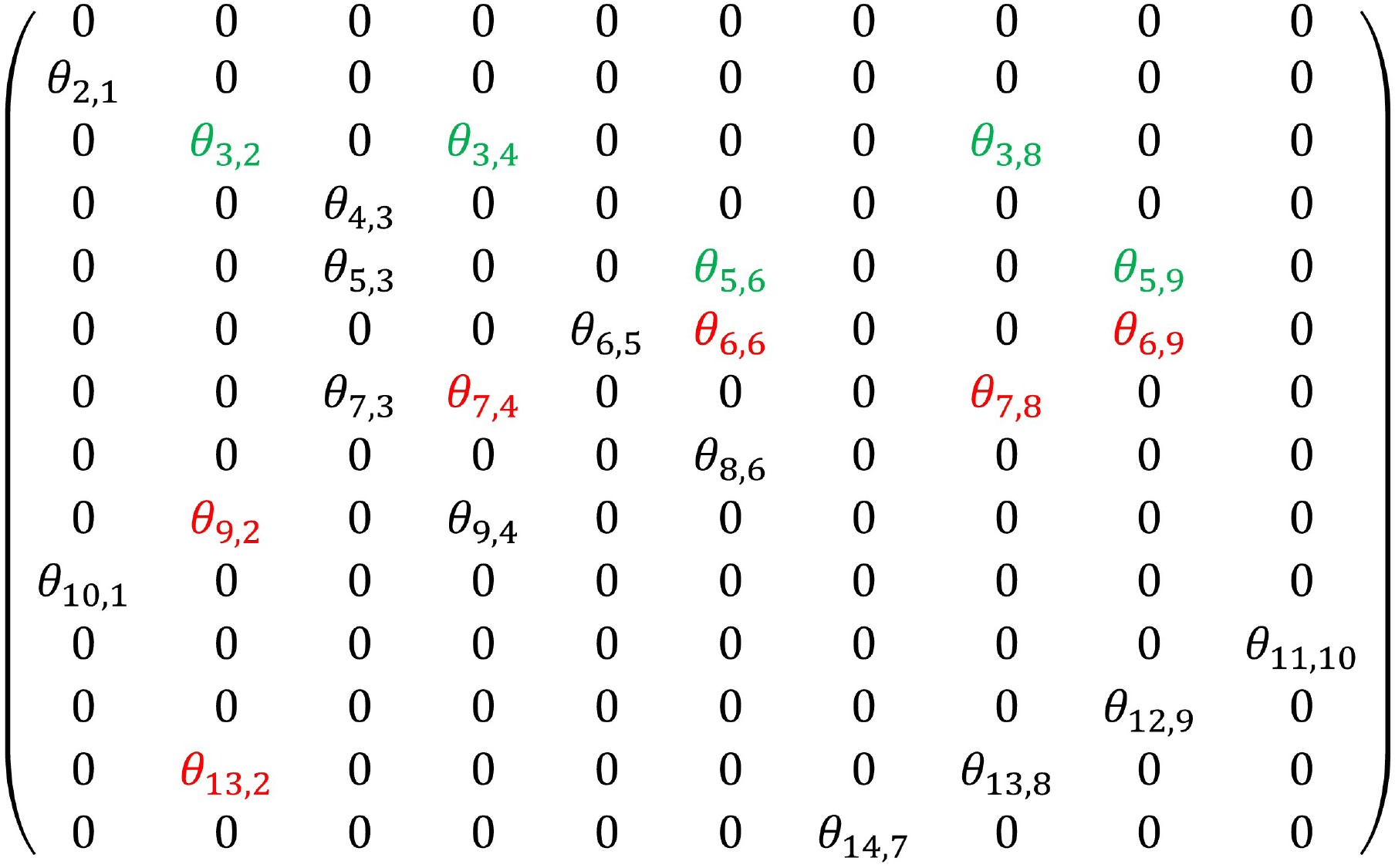
Θ matrix of the model. The matrix consists of 14 influenced fluxes (rows) and 10 influencing metabolites (columns). The matrix depicts all consistent elasticities (black, randomized between 0 and 1), all implemented activations (green, set to 1 or 0.33) and all implemented inhibitions (red, set between 0 and –0.99).

**Figure S3:**
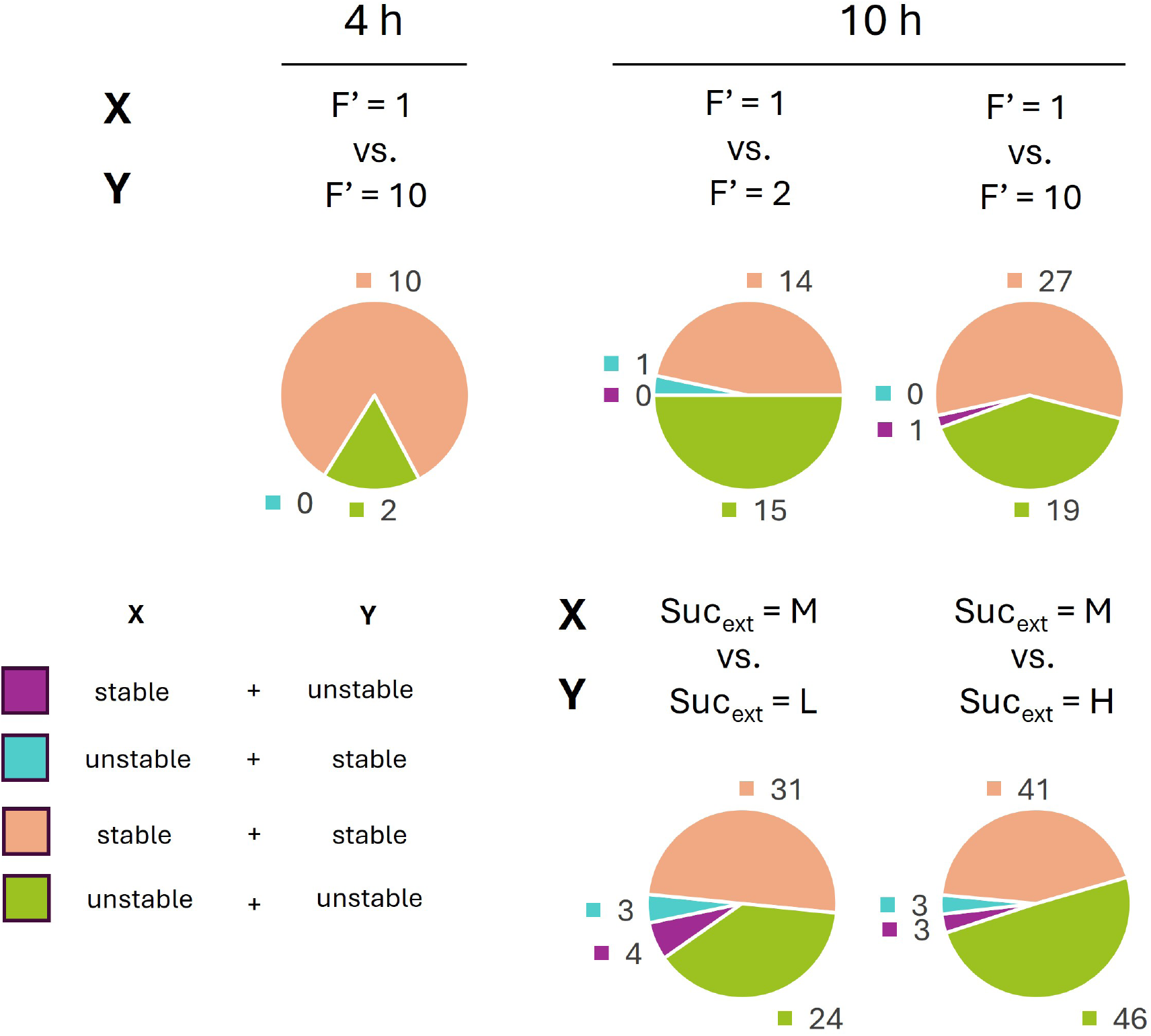
Stability analysis overview of different F flux sizes and external sucrose concentrations. A few of the 118 regulation combinations were tested with other fluxes F and other external sucrose concentrations for the 4 h datasets (A) and the 10 h datasets (B). The different tested external sucrose concentrations were 2/3 of the cytosolic sucrose concentration (L), the same as the cytosolic sucrose concentration (M) and 3/2 of the cytosolic sucrose concentration (H).

**Figure S4:**
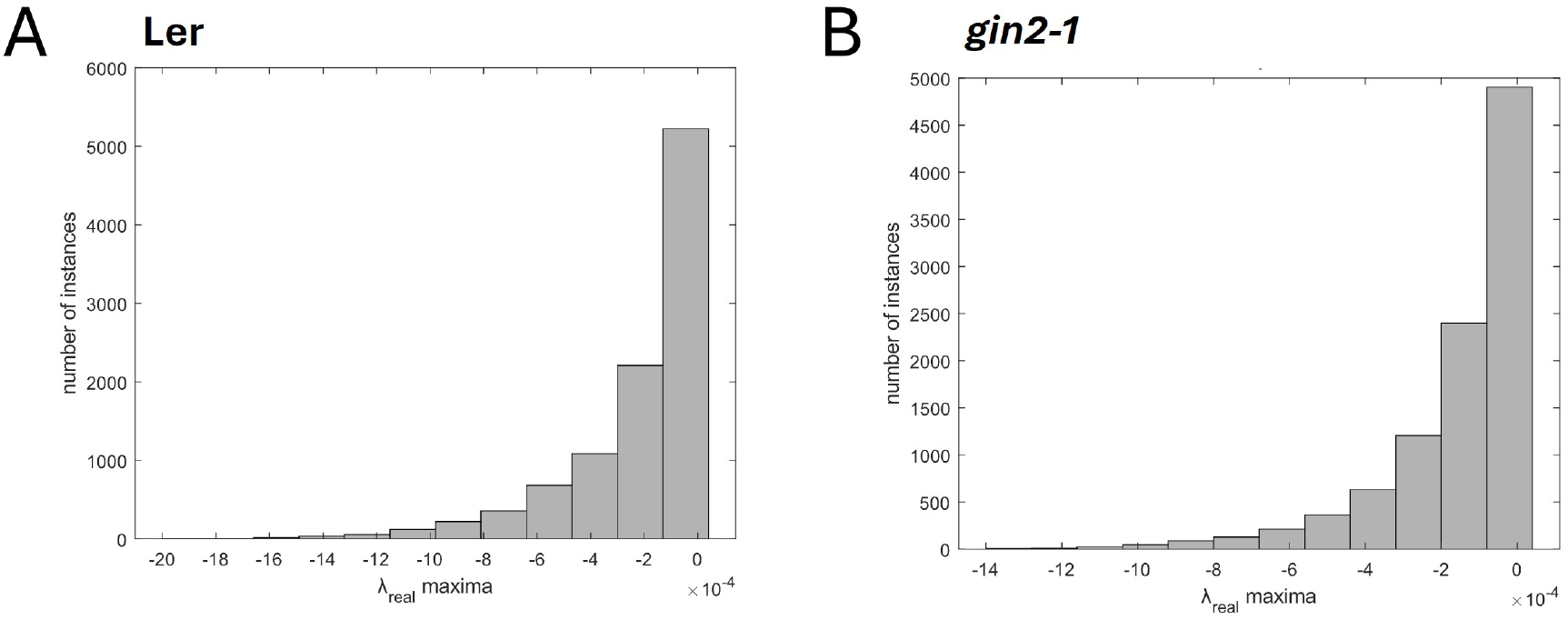
Histograms of Eigenvalue real part maxima. Histograms of maximal real part Eigen-values for 10^6^ calculations without regulatory instances with medium external sucrose concentration in (A) wild type *Arabidopsis thaliana* Ler and (B) mutant *gin2-1*.

**Figure S5:**
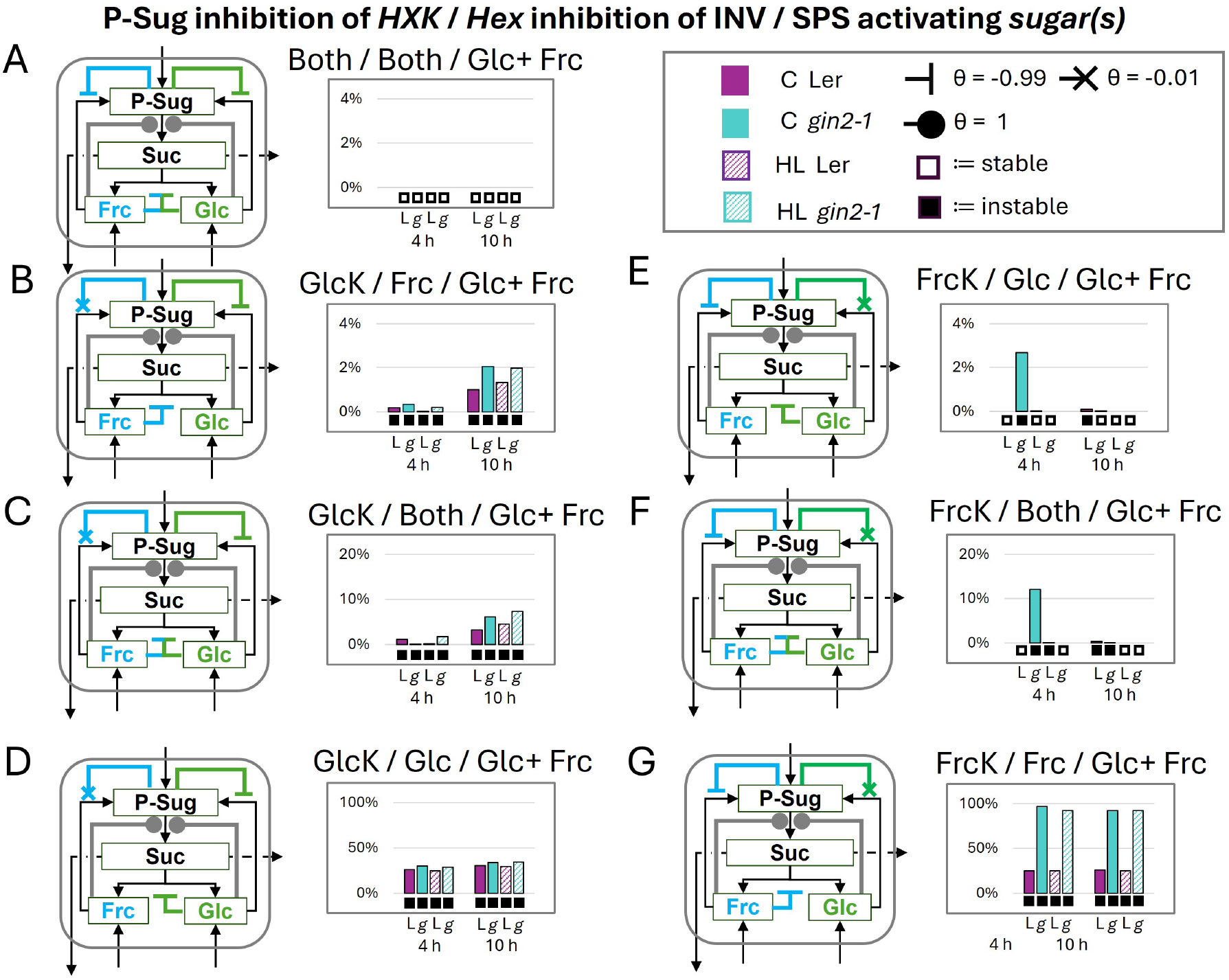
Influence of additional weak hexokinase inhibitions to prior regulations on the cytosolic sucrose cycle stability. Stability analysis of full activation of sucrose-phosphate synthase (SPS) by both non-phosphorylated hexoses. In addition to the prior regulations (Fig. 3) the noninhibited hexokinases were weakly (*θ* = –0.01) inhibited by P-Sug. The combinations are indicated above each diagram as follows: strong inhibition (*θ* = –0.99) of P-Sug on which *Hexokinase* / which *Hexose* inhibits cytosolic invertase / which *sugar(s)* activate the SPS. A) GlcK+FrcK/Frc+Glc (B – D) strong glucokinase inhibition (E –G) strong fructokinase inhibition. Note: Here only the cytosolic regulations are depicted, but the model consisted of the whole cell (Fig. 1). Boxes below diagrams indicate stability (empty box) and instability (black box). L:= Ler (purple, control/high-light:= filled/crosshatched), g:= *gin2-1* (turquoise, control (C)/high-light (HL):= filled/crosshatched), HXK:= hexokinase (GlcK:=glucokinase, FrcK:=fructokinase), INV:= invertase (here cytosolic), Frc:= fructose, Glc:= glucose, P-Sug:= phosphorylated sugars, Suc:= sucrose

**Figure S6:**
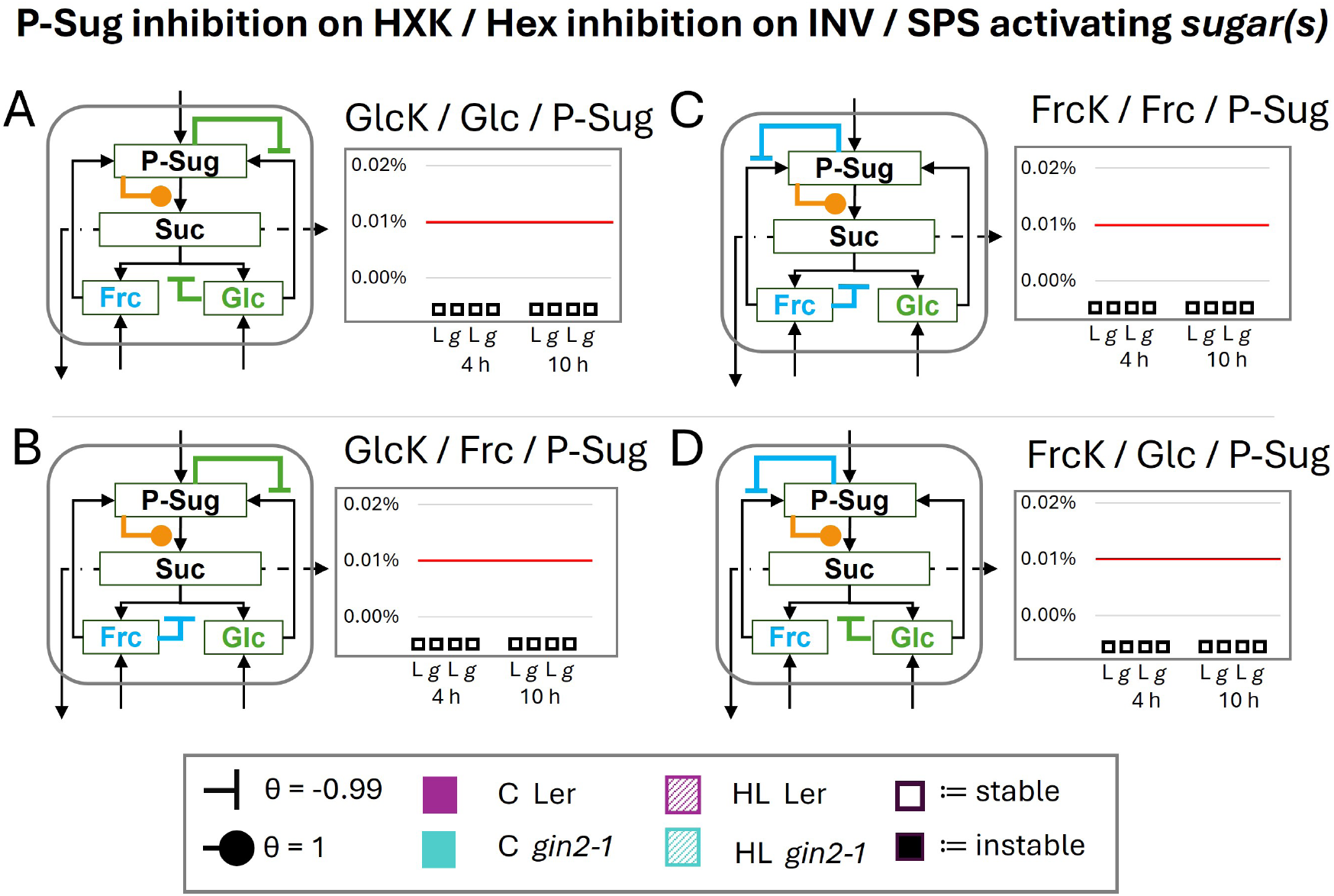
Influence of single activated sucrose-phosphate synthase (SPS) by phosphorylated sugars on the cytosolic sucrose cycle stability. Stability analysis of full activation of sucrose-phosphate synthase (SPS) by phosphorylated sugars (P-Sug). The subsequent combinations are indicated above each diagram as follows: inhibition of P-Sug on which *hexokinase* / which *hexose* inhibits cytosolic invertase / which *sugar(s)* activate the SPS. (A – B) glucokinase inhibition (C –D) fructokinase inhibition. Note: Here only the cytosolic regulations are depicted, but the model consisted of the whole cell (Fig. 1). Boxes below diagrams indicate stability (empty box) and instability (black box). L:= Ler (purple, control/high-light:= filled/crosshatched), g:= *gin2-1* (turquoise, control (C)/high-light (HL):= filled/crosshatched), HXK:= hexokinase (GlcK:=glucokinase, FrcK:=fructokinase), INV:= invertase (here cytosolic), Frc:= fructose, Glc:= glucose, P-Sug:= phosphorylated sugars, Suc:= sucrose; Red line cut-off for instability

## Notes

https://git.rwth-aachen.de/Lisa.Fuertauer/skm-sucrose-cycling

### Competing Interest Statement

The authors have declared no competing interest.

